# Metabolitin-based molecular drug delivery by targeting GPR158 in glioblastoma

**DOI:** 10.1101/2021.12.13.472376

**Authors:** Huashan Zhao, Wen Zhu, Jingwei Li, Jinju Lin, Xiaohua Lei, Pengfei Zhang, Jian V. Zhang

**Author notes:** These authors contributed equally to this work (Huashan Zhao, Wen Zhu). Correspondence should be addressed to Pengfei Zhang or Jian V. Zhang. or.

## Abstract

Glioblastoma multiforme (GBM) is a lethal form of intracranial tumor. One of the obstacles to treat GBM is the blood-brain barrier which limit the transportation of drugs into the tumor site. Here, based on our previous study on metabolitin (MTL) and osteocalcin, we generated a molecular drug delivery system that consisted of metabolitin and small molecules such as fluorescent dye or peptide drugs for diagnosis and treatment. And we designed a GBM diagnostic probe (MTL-ICG) and therapeutic peptide drug (MTL-NBD) that can cross the blood-brain barrier (BBB). In a NIR animal live imaging system, we found MTL-ICG can penetrate cross BBB and label the GBM site. The *in vitro* experiment showed that MTL-NBD had inhibitory effect on GBM cell line (U87-MG). Besides, after orthotopic transplantation of GBM into mouse cortex, treatment of MTL-NBD intravenously showed inhibition trend, which were similar with the effect of NBD, a known anti-tumor polypeptide drug. In addition, we found the GPR158, the receptor of osteocalcin, was also high expressed in grafting site. Taken together, these findings suggest that MTL is a promising cell penetrating peptide targeting GPR158 in GBM, which provide a novel delivery tool for GBM.

## Introduction

GBM is a tumor located in the central nervous system, with headache, vomiting, epilepsy as the main symptoms. According to the statistics of the American Central Brain Tumor Registry (CBTRUS), more than three out of every million people in the United States suffered from GBM from 2012 to 2016 (Ostrom et al., 2019). Primary GBM accounts for 95% of confirmed GBM cases; secondary GBM accounts for about 5% of confirmed cases (Ohgaki et al., 2004). Among them, the median survival of patients treated with chemotherapy alone was only 12.1 months, and that of patients treated with radiotherapy combined with TMZ was only 14.6 months (Stupp et al., 2005). To date, it is still a challenge to treat GBM, and there are big gap to understand BGM related signaling pathways, therapeutic drugs and treatment methods.

For GBM treatment drug, as the only standard treatment currently used in clinical practice, TMZ is a derivative of imidazoltetrazine, a precursor of the anticancer drug Temorda, which produces cytotoxicity through transformation into an active DNA alkylation agent (Denny et al., 1994). However, glioma IN-18, T98G and U138 cell lines have been reported to be consistently resistant to TMZ (Lee, 2016). It takes a long time to develop new drugs, and it is necessary to find effective drugs for GBM as soon as possible under the condition of poor prognosis and limited treatment for GBM. Therefore, some researchers have evaluated the effect of some marketed drugs on GBM. Studies suggest that psychotropic drugs, such as traditional antipsychotic chlorpromazine, are promising for the treatment of chemically-resistant glioma (Oliva et al., 2017). Atypical antipsychotic drug olanzapine can enhance the inhibitory effect of TMZ on GBM cell lines (Karpel-Massler et al., 2015). Tricyclic antidepressant promimazine can induce autophagy in GBM U87-MG cell lines (Jeon et al., 2011). Non-psychiatric drugs such as bendazole can promote the survival of glioma-bearing mice by inhibiting microtubule polymerization (De Witt et al., 2017). Clomiphene can inhibit IDH1 mutation of glioma cells, thus significantly inhibiting the growth of glioma in mice (Zheng et al., 2017). Although there are some achievements in the researches on GBM related pathways and drugs, there are not enough drugs with high efficiency, low toxicity and specific characteristics that can be used in clinical practice. So it is necessary to study other potential signaling pathways and drugs in the development of GBM.

In this study, we attempted to design a novel GBM probe and anti-GBM polypeptide, which consisted of transmembrane peptide MTL (YLGASVPSPDPLEPT), flexible link (GG) and ICG or NBD (TALDWSWLQTE). And we tested the loading capacity of the MTL *in vitro* and *in vivo*.

## Materials and Methods

### The experimental materials

The peptide MTL-NBD (YLGASVPSPDPLEPTGGTALDWSWLQTE) and positive peptide NBD (RQIKIWFQNRRMKWGGTALDWSWLQTE) (Friedmann-Morvinski et al., 2016) were synthesized by Shanghai Jiepeptide Biotechnology Co., LTD. Luciferase substrates sourced from GOLD BIO. DMEM, DPBS, 0.25% (W/V) trypsin solution, penicillin-streptomycin and fetal bovine serum (FBS) were sourced from Gibco Life Technologies (USA). Cell counting Kit (CCK8) was purchased from DOJINDO. OCT embedding materials are sourced from SAKURA Tissue-Tek.

### In vitro cell survival rate test

The effect of drugs on cell survival rate were tested by CCK-8 kit. First, peptide MTL-NBD was dissolved with DPBS and diluted to a series of different concentrations of 10 μM, 20 μM, 50 μM, 100 μM, 200 μM with DMEM, and peptide NBD was dissolved with 0.05% DMSO DMEM and diluted to 50 μM. U87 cells were cultured in a 25mL dish with DMEM containing 10% fetal bovine serum (FBS) and 1% penicillin-streptomycin at 5% CO_2_ 37 □. When the cells grew to 80% of the culture dish, the cells were digested with 0.25% trypsin and made into a single cell suspension. By cell counting, the cell suspension was diluted to 5×10^4^/ mL, with 100 μL adding into each well of the 96-well plate, 5000 cells per well. The cells were cultured for 24 hours to make the cells adhere to the wall. The 96-well plates were washed with DMEM for three times and replaced with fresh DMEM medium. They were starved for 4 hours. At 0 h, 24 h, 48 h, different concentration of MTL-NBD and 50 μM NBD were added into the cell culture medium. Fresh medium and drugs were replaced every 24 h. Adding CCK-8 detection solution into 96-well plate at 0 h, 24 h, 48 h and 72 h, with 10 μL per well, and incubating for 2 hours, then they were detected at 450 nm in detector. The cell survival rate (%) was equal to (drug group-A control group)/(control group-A blank group) ×100. The control group was added with PBS, and the blank group had no cells.

### In vivo model was established by orthotopic transplantation

Adult female C57 mice or BALB/C nude mice were used in experiment. The mice were purchased from Beijing Vital River Experimental Animal Technology Co., LTD. The mice were anesthetized by isoflurane and operated with isoflurane to maintain anesthesia. Eye ointment was applied to prevent eye desiccation. The mouse was fixed in the brain stereotatic injection instrument of small animals. After the skin was disinfected by povidone-iodine, a longitudinal incision about 1 cm was cut along the middle seam of the head with the sterilized surgical scissors. The point of intersection of sagittal and coronal fractures of the skull, namely Bregma point, was used as the origin. In experiment, a hole was drilled at x=−2.5 mm, Y =+1.5 mm, z=−3.5 mm and the needle was inserted at a uniform speed. After a 3-minute pause, the needle was raised 0.05mm and 1 μL U87 single cell suspension was injected, with a total of 1 million cells. The injection lasted for 5 minutes. After the injection, the needle was stopped for 5 minutes, and then the needle was slowly pulled out. The wound of the mouse skin was sutured and povidone-iodine was used for disinfection. The mice were observed until they woke up and put back into the cage.

### Histomorphological detection

For 5 days after brain injection, female C57 mice were sacrificed by neck dissection, and the entire brain was removed, immobilized with 4% PFA for 48 hours, and then dehydrated in 30% sucrose solution for 48 hours. After dehydration, the brains of mice were embedded in OCT and refrigerated at −80□ for more than 4 hours. The embedded brain tissue was taken out and balanced at −20□ for half an hour, sectioned in cryostat (Leica CML1950) with a thickness of 40μm. The sections were immunostained with GPR158 antibody and DAPI. The sections were observed under a microscope or a confocal microscope.

### In vivo detection under small animal live imaging system

It has been reported that the survival period of tumor-bearing nude mice could reach 28-29 days after injection of 1 million U87 cells into the brain of Balb/C nude mice (median survival period was 28.5 days) (Candolfi et al., 2007). In addition, U87 cells with luciferase were injected into the brain, and after injection of luciferase substrate, *in vivo* tumors can be imaged by IVIS system (Lai et al., 2019). For *in vivo* detection, mice were intraperitoneally injected with 150 μL luciferase substrate at a concentration of 15 ng/ml about 10 minutes before imaging. After isoflurane anesthesia, mice were placed into the system. The biotin luminescence imaging mode was selected. The aperture was 1, and the exposure time was 5 minutes.

Bioluminescence imaging was performed on IVIS system after brain injection in BALB/C nude mice. The mice were injected 20 mg/kg MTL-NBD polypeptide or 20 mg/kg NBD polypeptide intravenously for 4 consecutive days. The nude mice were imaged every two days, and tumor growth rate (tumor growth rate = tumor luminescence intensity after administration/tumor luminescence intensity before administration) was calculated based on the imaging results.

### Data Analysis

Statistical differences between data groups were tested by *t test,* and three or more groups were tested by ANOVA (GraphPad Prism 8). P<0.05 was considered as statistic significance.

## Results

### Probe design and Imaging of MTL-ICG in mouse brain

For detection purpose, we first design a probe (MTL-ICG) (Fig. 1A). By the live animal system test, we found the MTL could penetrate the BBB, compared with ICG only and PBS group (Fig. 1B), whereas the main organs of different groups showed similar pattern (Fig. 1C).

**Figure 1.**
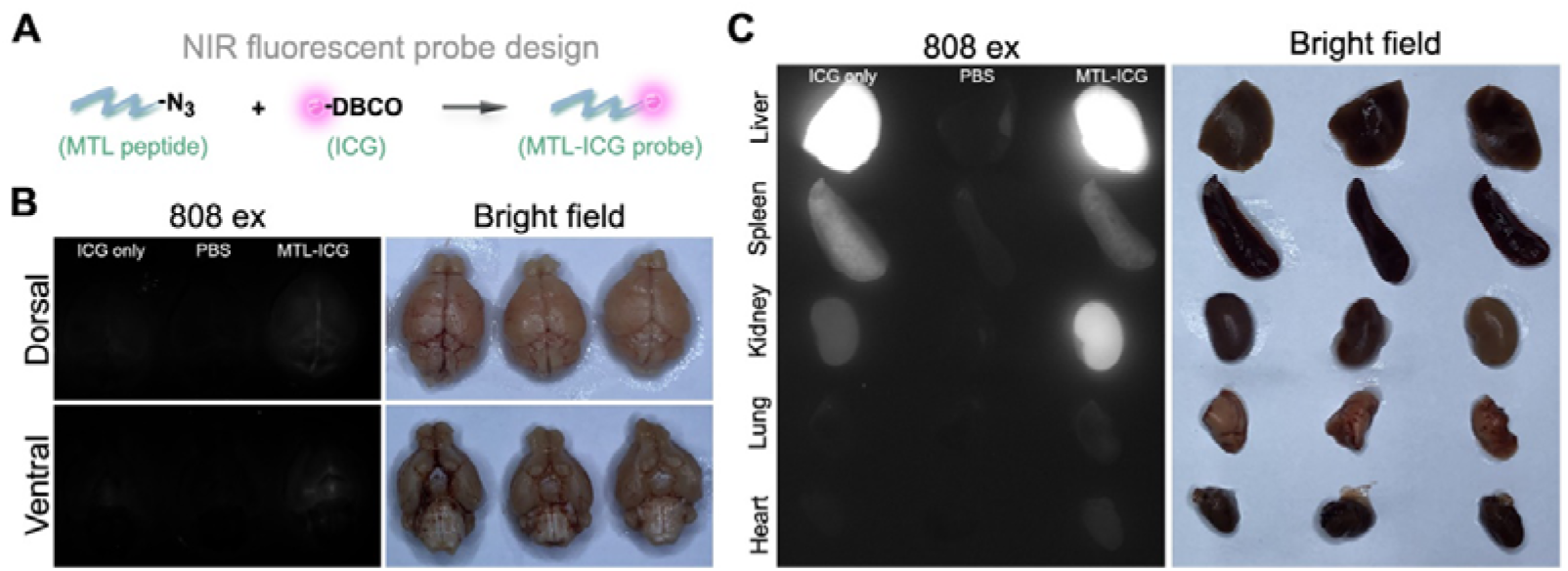
ICG probe design and *in vivo* cross BBB test. A) ICG probe design by click Chemistry. B) Live imaging of brain tissue after injection of the probe under 808 excitation light. C) The main organs imaging corresponding to panel B.

### Tumor imaging using MTL-ICG

The mice were sacrificed in day 5 after transplantation. Frozen sections were prepared for morphological observation. The tumor site was in the cerebral cortex of the mouse, and the U87 cells are clumped with clear boundary and stable morphological characteristics (Fig. 2A). The fluorescent signals could be detected by the live animal system, compared with scrambled peptide group (SCR), ICG only group and PBS group (Fig. 2B), and there were no obvious change in main organs (Fig. 2C).

**Figure 2.**
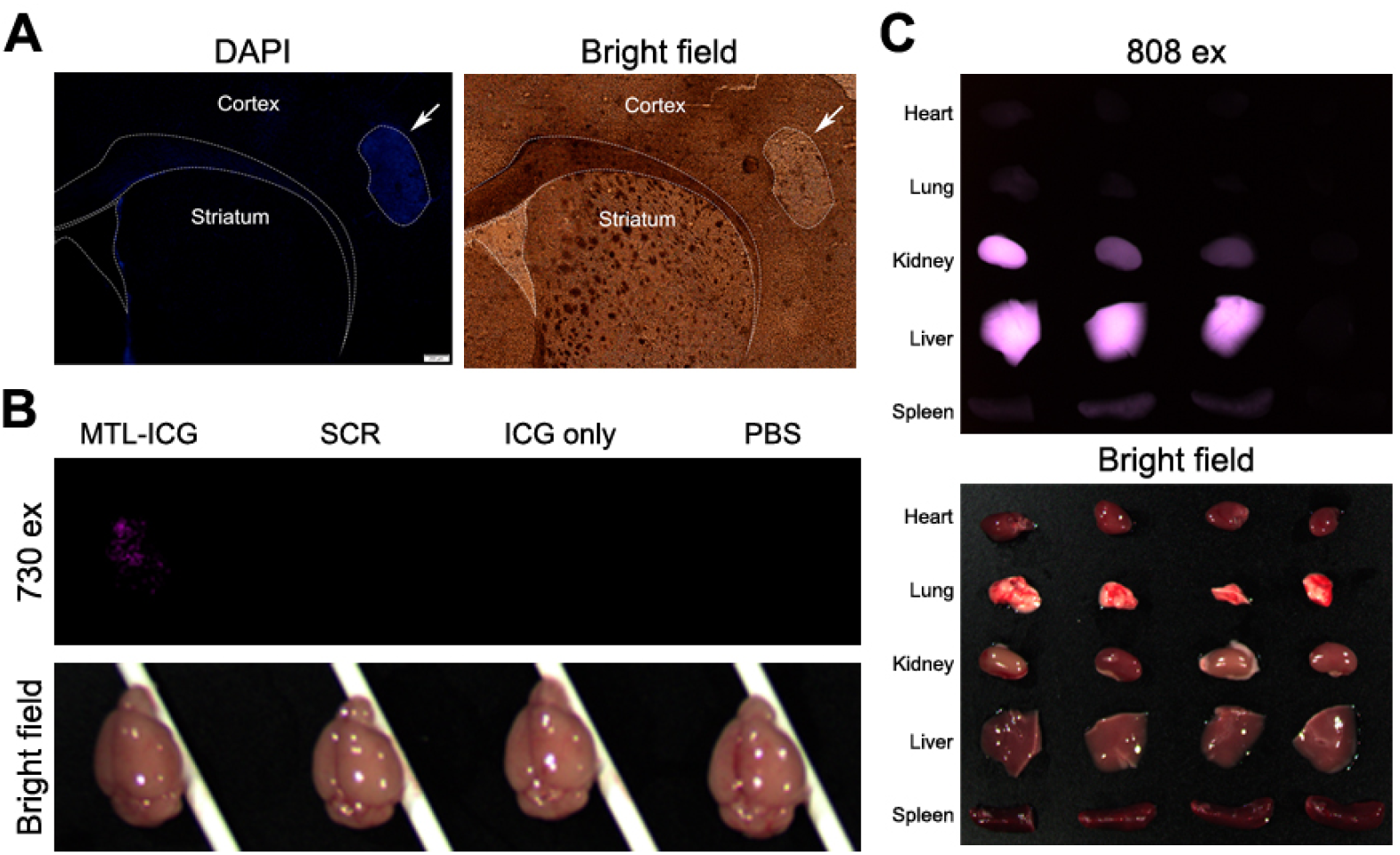
Histological morphology after cell stereotatic injection and probe treatment *in vivo*. A) Microscopic imaging results of mice brain after stereotactic brain injection, counterstaining with DAPI. B) The near infrared imaging of grafted brain, after the grafted mice were injected MTL-ICG, SCR, ICG only or PBS intraperitoneally for 24 hours. C) Main organ imaging of modeling mice brain corresponding to panel B.

### Design of MTL-NBD and in vitro test

U87 cells were co-incubated with MTL-NBD and NBD. The CCK-8 detection was performed to evaluate whether MTL-NBD had an inhibitory effect on GBM *in vitro*. The different concentrations of MTL-NBD can inhibit the growth of U87 cells, similar with the positive drug NBD (Fig. 3B).

**Figure 3.**
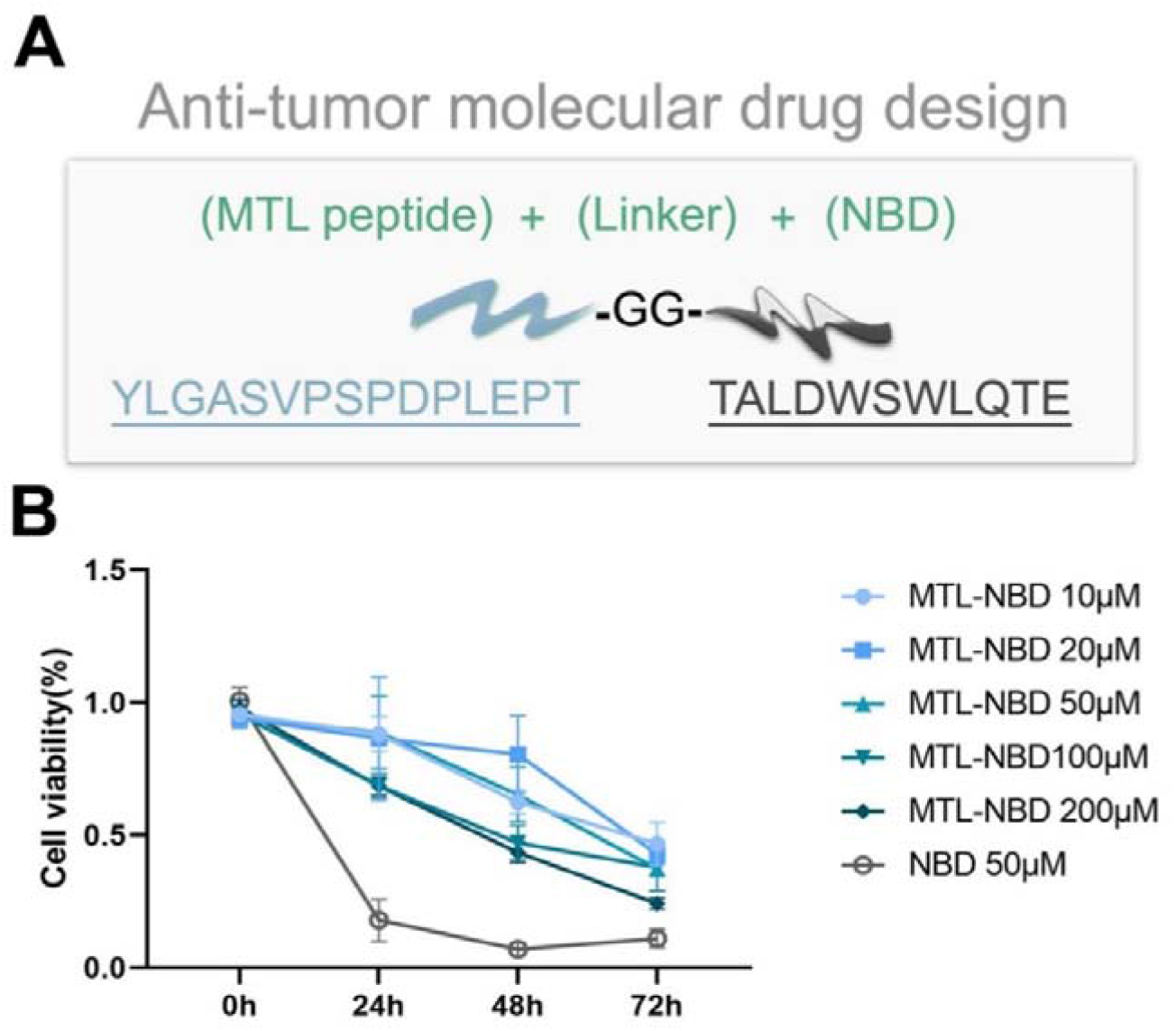
Design of MTL-NBD and inhibitory effect of MTL-NBD on U87-MG cell growth *in vitro*. A) Desing of peptide MTL-NBD; B) U87-mg cell lines were treated with different concentrations of NTL-NBD and NBD, and the cell survival rate was determined by CCK-8 assay kit. The data were the mean values of the three experiments, tested by two-way ANOVA, p<0.0001.

### Comparison of MTL-NBD and NBD in vivo

Twenty four hours after surgery, nude mice were injected with 20 mg/kg MTL-NBD or 20 mg/kg NBD intravenously every day, and *in vivo* imaging was performed in Day 1, Day 3 and Day 5 (Fig. 4A).. The luminescent intensity of the images was analyzed by software and tumor growth rate was calculated. According to luminescent intensity analysis, MTL-NBD and the positive drug NBD had certain similar inhibitory effect on the growth of brain GBM, with no significant difference (Fig.4B).

**Figure 4.**
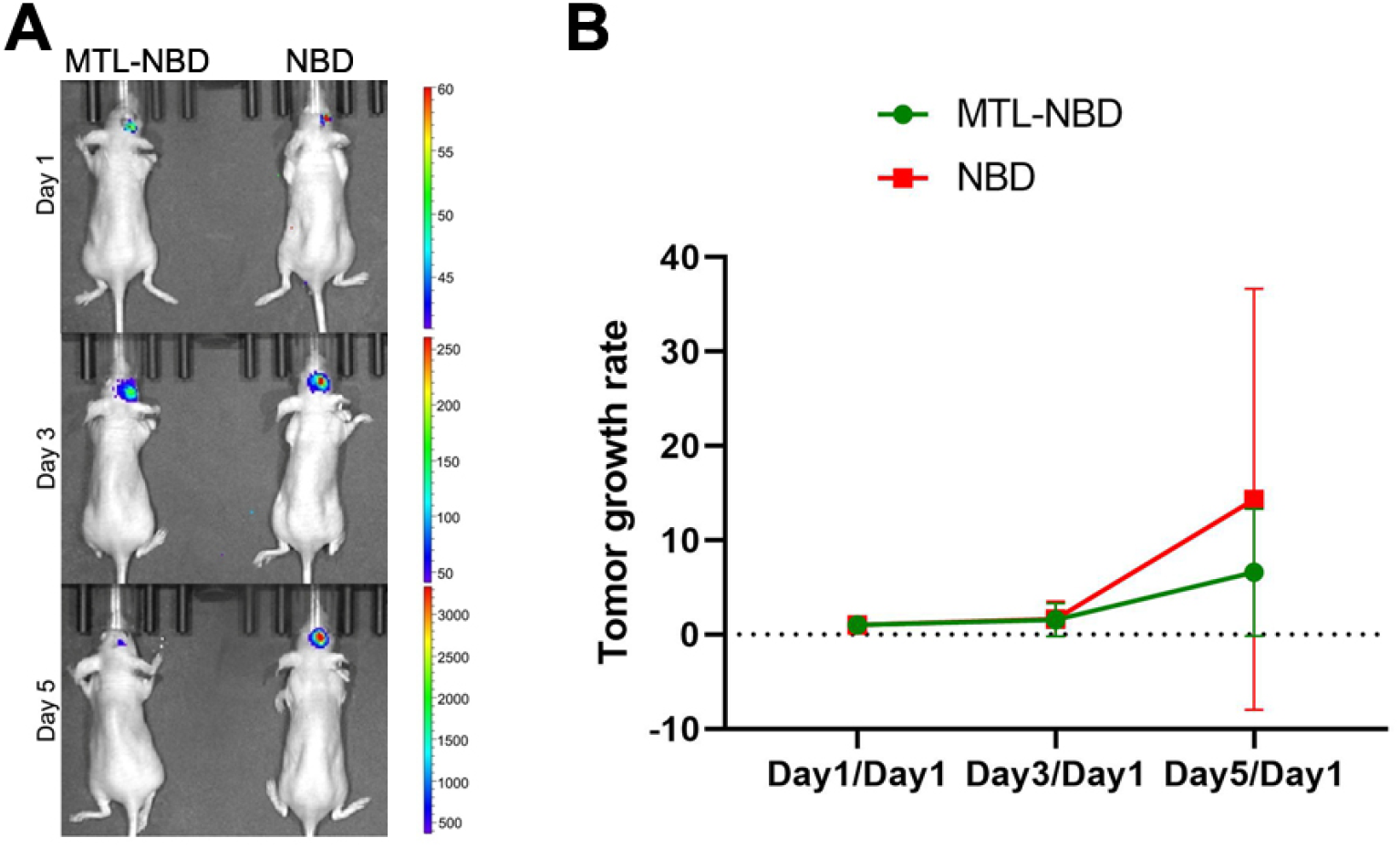
Inhibition effect of NTL-NBD on GBM growth *in vitro*. A) Animal live imaging results after stereotactic injection. B) Comparison of tumor growth rate, according to the luminescent intensity of tumor site.

### Expression of GPR158 in GBM graft

It has been reported the GPR158 is the receptor for osteocalcin, so we speculated MTL could bind to the GPR158 too. Thus, we tested the localization of GPR158 in the grafts, and we found GPR158 showed high expression in tumor sites (Fig. 5).

**Figure 5.**
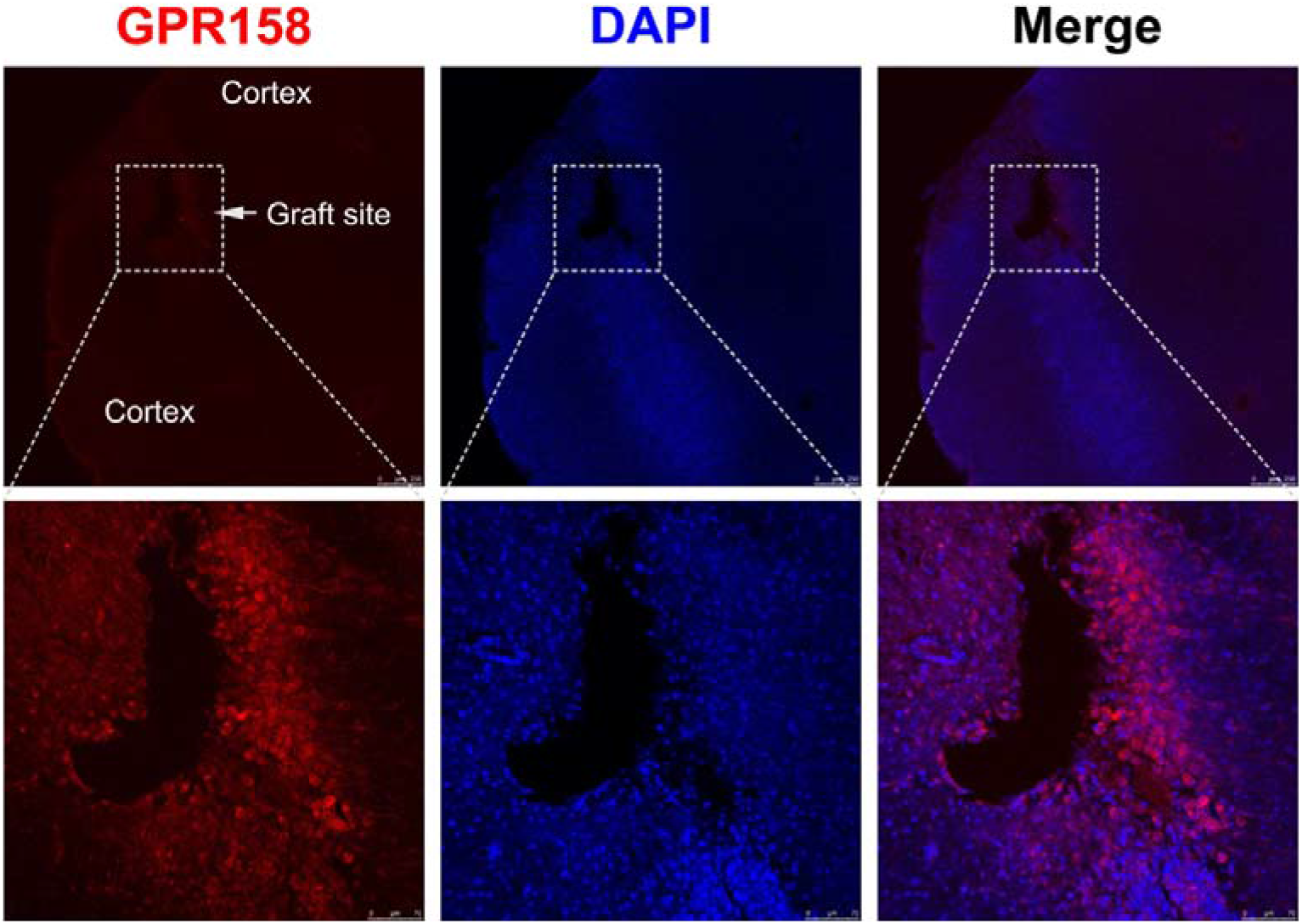
Detection of GPR158 in grafting site. GPR158 showed high expression in U87 transplanted area.

## Discussion

In this study, we demonstrated that MTL could be penetrate BBB and target GBM. -NBD had a significant inhibitory trend on GBM proliferation. The polypeptide NTL-NBD has the advantage for GBM therapy. Due to the composition of natural amino acids, NTL-NBD has low toxicity to animals. And it has good water solubility, namely 200 μL normal saline or DPBS can completely dissolve 1 mg NTL-NBD. The solvent would not introduce toxicity, so it can safely increase the dosage of NTL-NBD in animal experiments, with an optimistic application prospect. However, the water solubility of positive drug NBD is poor, so DMSO is needed to facilitate the dissolution, with potential toxicity, limiting the further application of NBD. In addition, 10% DMSO DPBS with 1000 μL as solvent could not completely dissolve 1mg of positive drug NBD, and only the mixture with loose precipitation could be obtained, which may affect the pharmacokinetics of NBD *in vivo* and reduce the concentration of NBD in tumor.

GBM is divided into isocitrate dehydrogenase-wildtype (GBM, IDH-wildType), isocitrate dehydrogenase-mutant (GBM, IDH-mutant), unshaped (GBM, NOS) and other rare types (Louis et al., 2016). Clinically, the frequency of IDH mutation in primary GBM is low, while the frequency of IDH mutations in secondary GBM is high (Balss et al., 2008). It has been reported that GBM may originate from neural stem cells (NSCs) in human subventricular zone (SVZ) tissues (Lee et al., 2018). After comprehensive analysis, the main pathway of GBM include receptor tyrosine kinase/RAS binding protein/intracellular phosphatidylinositol kinase (RTK, RAS/PI(3)K) signaling pathway, tumor suppressor protein (P53) signaling pathway and retinoblast gene (RB) signaling pathway (Network, 2013). According to our study, we GPR158 signaling might be involved in development of GBM.

In conclusion, *in vitro* and *in vivo* experiments have proved that the MTL-ICG and MTL-NBD we designed in this study could be used as probe and an inhibitory drug for GBM, indicating that MTL is a valuable peptide for the diagnosis and treatment of GBM, with a good application prospect as a new anti-GBM peptide drug. Next, the future work should figure out the relationship between MTL, GPR158 and GBM.

